# Risky home with easy food: Denning of a generalistic and widely distributed carnivore red fox

**DOI:** 10.1101/411686

**Authors:** Hussain S. Reshamwala, Neeraj Mahar, Rodolfo Dirzo, Bilal Habib

## Abstract

Dens are crucial for the survival of all canids, however, for a meso-carnivore like red fox, denning is of greater importance as they utilize dens all throughout the year for protection, resting and breeding. The red fox known for its generalist and opportunistic behavior and extremely good adaptability to the dynamic anthropogenic changes is the most widespread and successful wild land carnivore. With an ever-growing human population the choice of space for wild animals is limited and such adaptability is important for species survival. In this study we examined the denning preferences and den characteristics of red fox. Intensive surveys were conducted across two study sites Chiktan and Changthang. Foxes preferred to den on mountain slopes (43.70 ± 3.38 SE) where human disturbances were minimal. The red fox chose den sites closer to human settlements, water and road as compared to random points (Pillai’s trace = .487, F= 31.014 and P = 3.497e^-14^), which are risky and may expose the pups to humans but also provide the advantage of procuring anthropogenic food subsidies. In Changthang, foxes preferred to den at a greater distance (936 ± 223 meters) from human settlements as compared to Chiktan (403 ± 50 meters). The present research is of broad significance, given the increasing presence of human settlements within original animal ranges, even in remote harsh environments like the Trans-Himalaya.

## Introduction

Living in the Anthropocene, with an ever-growing human population, there is an increasing demand of land for agriculture, industries, housing and myriad other purposes. Land use change, including rapid urbanization has resulted in fragmentation of habitats available to wildlife across the world, forcing animals to live in close proximity to humans [1,2]. Living close to humans may have direct impacts on wildlife, such as poaching, legal hunting, isolation due to fencing or other barriers, road-kills, [3,4] or indirect impacts such as depletion of wild prey and provision of anthropogenic food subsidies [5]. Under such circumstances, carnivores present in human-dominated landscapes need to deploy numerous responses to thrive under those conditions. The red fox (*Vulpus vulpus*), known for its generalist and opportunistic behavior and extremely good adaptability to the dynamic anthropogenic changes [6], is the most widespread and successful wild land carnivore in the world [7]. Dens are crucial in the early development of many mammals, especially foxes as at this stage they are most vulnerable. While the use of dens maybe confined to the breeding period in canids like wolves, African wild dogs and dholes [8-10], foxes use their dens throughout the year [11]. Dens are utilized by the red fox either for resting during the day time, or for rearing the pups during the breeding season [12]. Availability of resources and shelter from predators are the main factors that determine den site selection [13]. Den site selection is extremely important to ensure pup survival. On the other hand, food resources originating from human activities like agricultural products, unmanaged waste and kitchen offal, are increasingly available and indeed present in the diet of animal populations [14]. Therefore, foxes are increasingly occurring in high densities near farmlands [15] and other urban and peri-urban areas, encouraging foxes to seek establishment in these human-dominated settings.

Denning close to humans occurs because of various reasons. In some cases human settlements not only provide opportunities to consume human-provided foods [16,14] but also enables them to avoid competition with meso-carnivores and reduce fear from apex predators (e.g., wolf and snow leopard), as these animals tend to avoid humans by selecting den sites away from human activity [17,18]. Some animals like skunks in USA and swift foxes in Canada have started exploiting manmade structures for making dens often in close proximity to roads and houses [19-21], others such as dholes and arctic foxes tend to live far from human settlements and therefore rarely interact with humans [9,22]. Although some human-dominated landscapes seem to provide both resources and protection from predators, some species (e.g., wolves) typically den away from human disturbance often shifting dens to avoid humans during the early life of their young [23]. Foxes in particular tend to have breeding dens in man-made structures and human dominated landscapes and they are known to shift dens when disturbed [24].

In some regions of the world, human culture and religion also have direct or indirect roles in determining the occurrence of wildlife species. Black Kites in Delhi, for example, choose nesting sites closer to Muslim communities because of their ritualized feeding practices and high density of human activity leading to large accumulations of food waste [25]. Reshamwala et al. [26] also found higher red fox encounter rates in areas with Muslim communities as these areas provided more anthropogenic food subsidy and also favoured fox communities via dog elimination.

Previous research on fox diet composition and distribution in the Trans-Himalayan region showed a strong influence of anthropogenic food subsidies on the pattern of red fox diet and occurrence across this region^26^. The Trans-Himalayan region harbors a highly inhospitable cold desert biotope with a short summer and Arctic-like winters [27]. Arid ecosystems, including the Trans-Himalayas, are characterised by low ecosystem productivity and in turn low animal densities [28]. The Indian trans-Himalayan region is also experiencing a rapid change in land-use pattern due to urbanization and increasing human population [29], with increasing incursion of human encroachments into the ranges of wild animals, including foxes.

In this study we build on our previous work on diet changes in foxes and turn to examine denning preferences and den characteristics of red fox. The availability and use of denning sites are important aspects in the ecology of most canids and indicative of breeding units within the habitat [24]. Knowledge about den ecology is crucial for understanding the denning strategies of targeted species in different environments, such as the Trans-Himalayan biome, ultimately leading to their reproductive success [30, 9, 31]. Given the increasing omnipresence of human settlements within original animal ranges, even in remote, harsh environments like the Trans-Himalaya, make this present research of broad significance.

## Materials and Methods

### Study Site

The Indian Trans-Himalayan landscape ranges from the northern border of the country and extends upto Tibet (China) and Nepal. The trans-Himalayan region of Ladakh comprises of high altitude dry rugged mountains with an average altitude varying from 2800 to 7000 meters above sea level and precipitation of about 100mm [32] which is majorly in the form of snow. The climate is cold and arid with temperatures dropping down to −30°C in winters, providing a short growing season for plants from May to August [33]. The region is sparsely populated and clustered with a density of about 4.9 people/km^2^ and most of the people are locally engaged in traditional pastoralism and agro pastoralism, although sometimes they also work as trekking guides and engage in tourism associated activities [34, 35]. The vegetation is very sparse with few alpine meadows dominated by *Kobresia* spp., *Carex* spp., *Potentilla* spp., and *Nepeta* spp. and shrublands dominated by *Hippophae* spp., *Salix* spp., and *Myricaria* spp. [36]. Compared to India’s other terrestrial ecosystems, where most extant wildlife populations survive inside protected areas, in the Himalayan and Trans-Himalayan landscapes wildlife populations are not restricted to protected areas, but occur across the landscape [37]. Snow leopard (*Panthera uncia*), wolf (*Canis lupus chanco*), lynx (*Lynx lynx*), pallas’s cat (*Octolobus manul*), mountain weasel (*Mustela altaica*), stoat (*Mustela* spp.) and stone marten (*Martes* spp.) are the carnivores found in this area. Voles (*Alticola* spp.), pikas (*Ochotona* spp.), hares (*Lepus* spp.) and marmots (*Marmota* spp.) are important prey species for the red fox found in this area.

We conducted intensive sampling in two representative sites across the landscape-Chiktan and Changthang (Fig. 1). Chiktan is densely populated and dominated by Muslim community. There is high anthropogenic food subsidy which governs the high fox occurrence here [26]. Changthang has a comparatively less rugged, flat terrain and is predominated by the Buddhist community. While Chiktan is completely devoid of dogs, Changthang has a good population of dogs.

**Figure 1.**
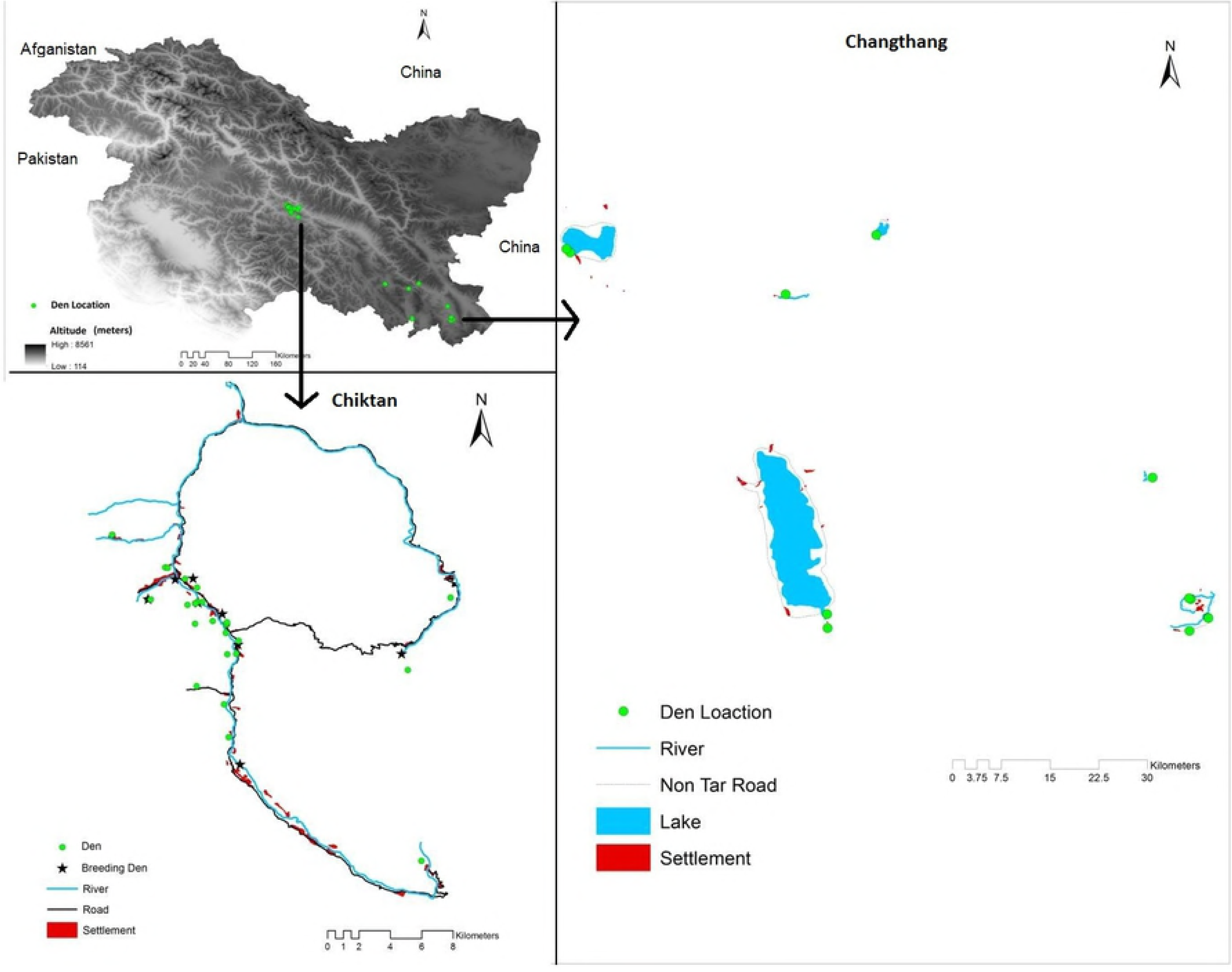
Den sites of red fox across the study sites of Chiktan and Changthang in the Trans-Himalayan region of Ladakh, India.

### Data collection and Analyses

Fox dens were intensely searched in the area of Chiktan from June 2015 to June 2017 using information obtained from sign surveys as well as secondary information from villagers and shepherds. Dens were searched from the Changthang region in 2016. A total effort of 814.5 km (638.47 km in Chiktan and 142.65 km in Changthang) was put in to locate the dens. Dens were identified as fox dens if they showed signs such as scats, tracks, occasional sightings, hair, bones, hides, feather or other prey remains outside or inside dens. Seven foxes were also radio-collared. Active dens once located were monitored regularly. The maximum length width and depth were measured when possible. Breeding dens were identified by sightings of pups and their scat and pug marks. These dens also had greater number of scats (adult and pups >50), bones, animal hides and often shoes, cloth, etc. around them. Den parameters were noted such as distance to water, road, house and cover using a range finder. Slope and aspect were also measured using a clinometer and compass. Aspect was classified into three classes sunny (112.5°-202.5°), half-sunny (22.5°-112.5° and 202.5°-292.5°), and shady (292.5°-22.5°) [38]. We generated 51 random points in ArcGIS version 10.1 and compared them to the den sites with the variables distance to road, distance to the nearest human settlement and distance to water. Multivariate analysis of variance (MANOVA) test was done to find out their denning preferences with random points as well as across the two different study sites. All the analysis was done in SPSS V 17.0 and R statistics (R Studio).

### Ethical statement

The radio-collaring of red fox was done with the permission of Department of Wildlife Protection, Jammu & Kashmir vide number (WLP/Res/2016/51-56) and no foxes were harmed/injured. The study was in accordance with the approved guidelines of Jammu & Kashmir Wildlife Protection Act (1978).

## Results

### Den Characteristics

Most of the dens (56.86%) were found in naturally occurring rocky crevices (Table 1). Foxes also dug dens in sandy substrate (37.25%). A few dens were also found in slate stones (5.88%). Foxes preferred to den on mountain slopes (43.70 ± 3.38 average) where human disturbances were minimal. Dens were also found in seabuck thorn scrub (*Hippophae* spp.) near agricultural crops from the study site of Chiktan, but because of the impenetrable dense scrub, few of these could be located (n=2). Majority of the dens used as a temporary shelter had single entrance (n=28) and double entrance (8). The breeding dens had three or more entrances (n=8) as alternative routes for the pups to escape when in danger. Since most of the dens were in naturally occurring rocky crevices and on steep rugged terrain, dimensions of only few dens could be noted (n=29). The den openings had an average length of 57.18 ± 7.59 cm and a width of 32.70 ± 2.57 cm. The opening of the dens was like a keyhole and the depth of only a few dens could be measured (n=13). The average depth of dens was 124.92 ± 23 cm.

**Table 1.** Fox den characteristics in Trans-Himalaya, India.

| Den Characteristics |  |  |  |  |
| --- | --- | --- | --- | --- |
| | N (Dens) | Minimum | Maximum | Mean ± (Std.error) |
| Length (cm) | 29 | 20.00 | 157.00 | 57.18 ± 7.59 |
| Width (cm) | 29 | 14.00 | 72.00 | 32.70 ± 2.57 |
| Depth (cm) | 13 | 40.00 | 283.00 | 124.92 ± 23.00 |
| Number of openings | 51 | 1 | 5 | 1.88 ± .162 |
| Slope | 51 | 0 | 76 | 43.70 ± 3.38 |
| Substrate | 51 | Sandy % | Rocky % | Slate % |
|  |  | 37.25 | 56.86 | 5.88 |
| Aspect | 51 | Sunny % | Half-sunny % | Shady % |
|  |  | 23.52 | 49.02 | 27.45 |

### Denning preferences

The red fox showed significant denning preferences with respect to their distances from human settlements, water and road as compared to random points (Pillai’s trace = .487, F= 31.014 and P = 3.497e^-14^) (Fig 2). Both the study sites, Chiktan and Changthang showed significant difference (Pillai’s Trace = .211, F= 4.194 and P = .010, Supporting information 1) in the denning parameters, but this was due to the combined effect of all three parameters. There was significant difference in the study sites only with respect to distance to human settlements (P =.001), while the difference between distance to water and road across the two study sites remained insignificant (P =.119 and P= .374).

**Fig 2.**
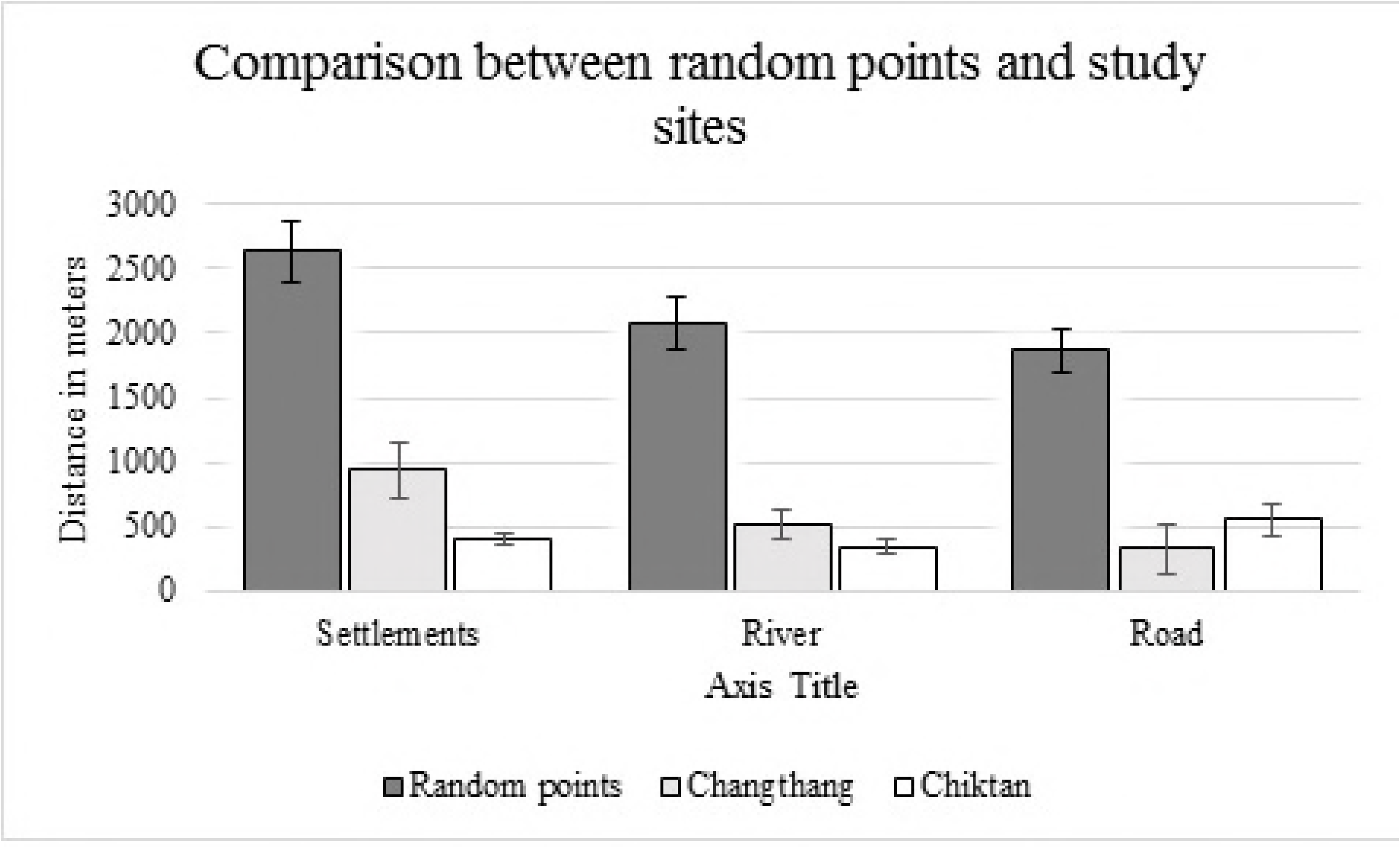
Comparison of dens and random points with respect to human settlements, rivers and roads in two different study sites.

In Chiktan 74.35 % (n=39) of the dens were located within 500 meters from human settlements whereas beyond 500 meters only 25.64 % of the dens occurred (Fig 3). Changthang had most of the dens i.e. % (n=12) located beyond 500 meters of human settlements. The optimum distance for denning for fox from human settlements was 201-300 meters in Chiktan and 401-500 meters for Changthang.

**Figure 3.**
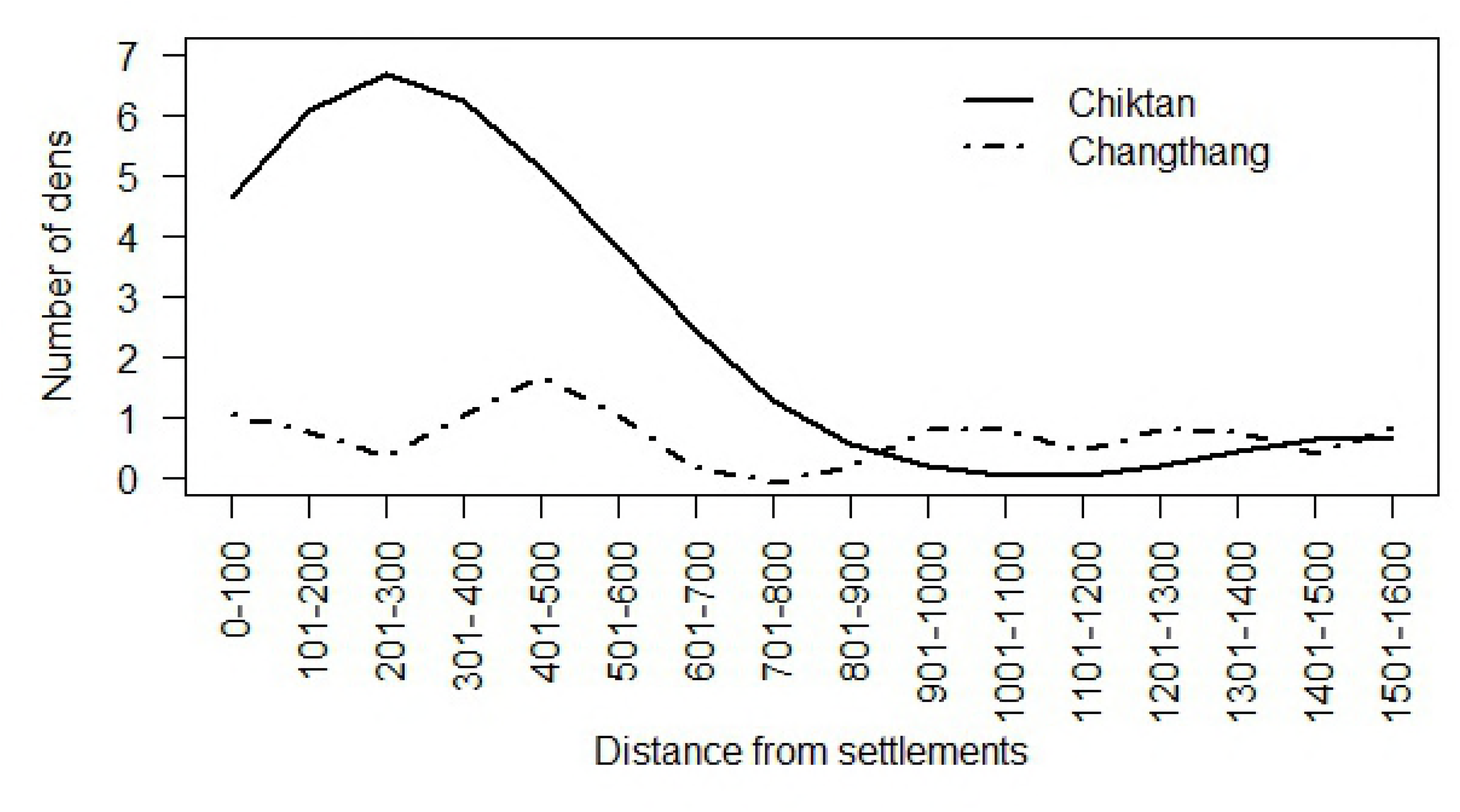
Distance of fox dens with respect to human settlements at two study sites in Trans-Himalaya, India.

## Discussion

Breeding dens usually had two to five entrances as escape routes for pups when approached by humans or predators. Multiple den-openings for raising their offsprings are also reported in other fox species [39-41]. Kilgore [11] also suggested large number of openings may indicate long term residency. It was observed by Manakadan et al. [42] that often small rodents may act as den providers for the Indian fox, which may expand an existing network of rodent burrows to reduce energetic costs of digging dens. In the present study, we found that foxes mostly utilized (n=32) naturally occurring rocky crevices and even utilized man-made structures like graves to reduce the energetic cost of digging and dug dens only when no such structures were available (n=19). Stanley [43] and Young and Jackson [44] reported that red foxes and coyotes *(Canis latrans)*, use dens only in severe winter weather and in the breeding season, however in this study we found that foxes used multiple dens throughout the year. Dens facing southern slopes in northern hemisphere have higher soil temperature [45]. A study by Chesemore [46] also suggested warmer microclimate inside dens having a southern orientation. Studies on red fox [38] and kit fox [41] have shown that they select southerly aspects for denning. Half-sunny aspect seems to provide moderate temperature conditions and most of the dens (49.02%) were found in this aspect. However, the half-sunny aspect also covers a wider angle i.e. north, north-west, north-east and east and thereby maybe biased. Studies by Zagrebel’nyi [47] on arctic fox suggested no significant correlation of aspect with their denning. In the current study also aspect might not be factor for selecting den sites. Foxes living in arid landscapes may select dens near permanent water bodies [48, 49].

Red foxes denned in a variety of situations, but did show certain preferences. They preferred sites closer to water bodies, roads and human habitation than random points, perhaps due to increased anthropogenic food subsidies. Although the data for prey density was not estimated, the foxes chose dens closest to water bodies in both the study sites perhaps because of good rodent population near these sites as also suggested by Szor et al. [30]. One reason to den near roads and human settlements could be to avoid other carnivores like wolf and snow leopard that are not as tolerant of humans. Similar aversion has also been observed in red foxes of North Dakota and east-central Illinois, USA where they preferred to choose denning sites closer to roads to avoid coyotes [50]. Another possible benefit to den near the roads could be to opportunistically scavenge on road-killed reptiles, invertebrates, and rodents. The foxes may do so during the night, as there are almost no vehicle plying on these roads at night. The low availability of natural resources in this landscape may also lead foxes to seek den sites close to human habitation for food resources.

The dens of Changthang primarily differed from dens of Chiktan with regard to their distance from human settlements. In Changthang the foxes preferred to den at a greater distance from the human settlements (936 ± 223 meters) as compared to Chiktan (403 ± 50 meters). While food subsidies are available at both the study sites near human settlements, the main reason why foxes choose to den away from the food resource in Changthang could be the presence of dogs. The presence of dogs is further determined by the local community in our study sites. Chiktan comprises of Muslim community which discourage the keeping of dogs and are highly intolerant to other top predators like wolves and snow leopards as well, while Changthang mainly comprises of the Buddhists community who are not only tolerant but also own dogs for the protection of their livestock [51]. Dogs are known to harass red foxes and compete with them for resources [52]. Reshamwala et al. [26] also found that the consumption of anthropogenic food subsidies in the red fox diet decreased in areas where dogs are present. The foxes also made dens closer to roads in Changthang (326 ± 198 meters) as compared to Chiktan (552 ± 124 meters). This could be attributed to the fact that there are very few vehicles in Changthang which ply on non-tar roads and dirt trails. Hence, road does not seem to be a disturbance factor in Changthang, whereas roads in Chiktan are comparatively busy with both people and traffic.

## Conclusion

To den in proximity of human settlements seems to be like two sides of a coin. On one side, this provides easy access to anthropogenic food subsidies without facing competition from other carnivores, while on the other, this exposes the fox and their pups to humans which can be potentially harmful. As a result of predation on poultry and livestock people often smoke dens (n=2) and kill pups of fox and wolves in the study site. In spite of the high mortality of pups induced by humans, the ability to exploit humans as sources of food and shelter appears to be a behavioral adaptation that helps foxes to survive in the arid Trans Himalayan landscape. In the present era of anthropocene where human culture and religion, management policies and other anthropogenic factors primarily govern the presence and distribution of wildlife species; the effective conservation of carnivores by protecting these denning sites beyond the boundaries of protected areas such as the human-dominated agricultural landscapes is of paramount importance [53]. Further work involving other areas of this landscape should be conducted to identify variations in denning ecology and movement patterns of this species in the Trans Himalayan region. A long-term study on den ecology could also provide good estimates of reproductive success and survival of the red fox.

## Supporting Information

**S1** Table. Mutlivariate analysis of variance (MANOVA) test results of red fox den site selection in Chiktan and Changthang

## Acknowledgements

This research was supported by Wildlife Institute of India, Grant-in Aid. We are thankful to Director and Dean, Wildlife Institute of India, for support and encouragement. We are grateful to the Department of Wildlife Protection, Govt. Jammu & Kashmir for providing the necessary permissions. We are thankful to Jigmet Takpa, CCF Leh and Samina A. Charoo, Research officer DWLP, J&K for institutional support and encouragement. Ninad Mungi is acknowledged for his valuable comments. Mr. Zakariya and Mr. Rigzin are acknowledged for helping out in the field work. First author also acknowledges fellowship from National Mission on Himalayan Studies and Idea Wild Grant for providing the necessary field equipment.

## Author contributions

The data collection and statistical analyses was done by H.S.R and N.M, and B.H., R.D. supervised the work and co-wrote the manuscript.

